# BrainGNN: Interpretable Brain Graph Neural Network for fMRI Analysis

**DOI:** 10.1101/2020.05.16.100057

**Authors:** Xiaoxiao Li, Yuan Zhou, Nicha Dvornek, Muhan Zhang, Siyuan Gao, Juntang Zhuang, Dustin Scheinost, Lawrence Staib, Pamela Ventola, James Duncan

**Author notes:** Corresponding Author: Xiaoxiao Li. Equal contribution (Nicha Dvornek, Muhan Zhang, Siyuan Gao).

## Abstract

Understanding which brain regions are related to a specific neurological disorder or cognitive stimuli has been an important area of neuroimaging research. We propose BrainGNN, a graph neural network (GNN) framework to analyze functional magnetic resonance images (fMRI) and discover neurological biomarkers. Considering the special property of brain graphs, we design novel ROI-aware graph convolutional (Ra-GConv) layers that leverage the topological and functional information of fMRI. Motivated by the need for transparency in medical image analysis, our BrainGNN contains ROI-selection pooling layers (R-pool) that highlight salient ROIs (nodes in the graph), so that we can infer which ROIs are important for prediction. Furthermore, we propose regularization terms—unit loss, topK pooling (TPK) loss and group-level consistency (GLC) loss—on pooling results to encourage reasonable ROI-selection and provide flexibility to encourage either fully individual- or patterns that agree with group-level data. We apply the BrainGNN framework on two independent fMRI datasets: an Autism Spectrum Disorder (ASD) fMRI dataset and data from the Human Connectome Project (HCP) 900 Subject Release. We investigate different choices of the hyper-parameters and show that BrainGNN outperforms the alternative fMRI image analysis methods in terms of four different evaluation metrics. The obtained community clustering and salient ROI detection results show a high correspondence with the previous neuroimaging-derived evidence of biomarkers for ASD and specific task states decoded for HCP. We will make BrainGNN codes public available after acceptance.

## 1 Introduction

The brain is an exceptionally complex system and understanding its functional organization is the goal of modern neuroscience. Using fMRI, large strides in understanding this organization have been made by modeling the brain as a graph—a mathematical construct describing the connections or interactions (i.e. edges) between different discrete objects (i.e. nodes). To create these graphs, nodes are defined as brain regions of interest (ROIs) and edges are defined as the functional connectivity between those ROIs, computed as the pairwise correlations of functional magnetic resonance imaging (fMRI) time series, as illustrated in Fig. 1.

**Fig. 1:**
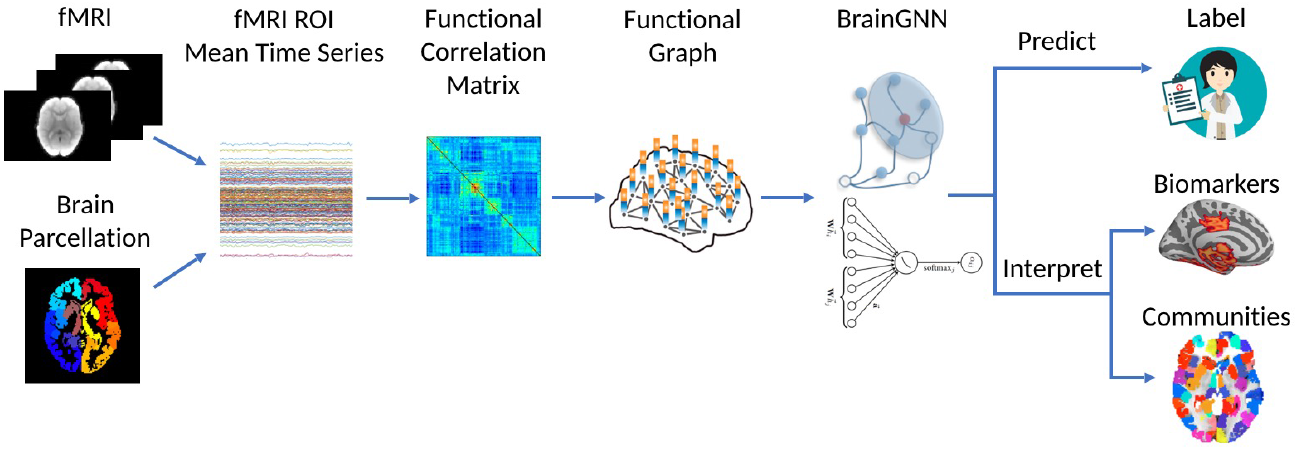
The overview of the pipeline. fMRI images are parcellated by an atlas and transferred to graphs. Then, the graphs are sent to our proposed BrainGNN, which gives the prediction of specific tasks. Jointly, BrainGNN selects salient brain regions that are informative to the prediction task and clusters brain regions into prediction-related communities.

Traditional graph-based analyses for fMRI have focused on two-stage methods: stage 1—feature engineering from graphs—and stage 2—analysis on the extracted features. For feature engineering, studies have used graph theoretical metrics to summarize the functional connectivity for each node into statistical measurements [58,32]. Additionally, due to the high dimensionality of fMRI data, usually ROIs are clustered into highly connected communities to reduce dimensionality [44,15] or perform data-driven feature selection [52]. For these two-stage methods, if the results from the first stage are not reliable, significant errors can be induced in the second stage.

The past few years have seen growing prevalence of using graph neural networks (GNN) for end-to-end graph learning applications. GNNs are the state-of-the-art deep learning methods for most graph-structured data analysis problems. They combine node features, edge features, and graph structure by using a neural network to embed node information and pass information through edges in the graph. As such, they can be viewed as a generalization of the traditional convolutional neural networks (CNN) for images. Due to their superior performance and interpretability, GNNs have become a widely applied graph analysis method [35,34,61,62,24,45]. Most existing GNNs are built on graphs that do not have a correspondence between the nodes of different instances, such as social networks and protein networks. These methods—including the current GNN methods for fMRI analysis—use the same embedding over different nodes, which implicitly assumes brain graphs are translation invariant and nodes on brain graphs (brain ROIs) are identical. However, nodes in the same brain graph have distinct locations and unique identities. Thus, applying the same embedding over all nodes is problematic. In addition, although recent studies have investigated group-level [38,56,50,61] and individual-level [8,41,37] neurological biomarkers, few GNN studies have explored both individual-level and group-level explanations, which are critical in neuroimaging research.

In this work, we propose a graph neural network-based framework for mapping regional and cross-regional functional activation patterns for classification tasks, such as classifying neurodisorder patients versus healthy control (HC) subjects and performing cognitive task decoding. Unlike the existing work mentioned above, we tackle the limitations of considering graph nodes (brain ROIs) as identical by proposing a novel clustering-based embedding method in the graph convolutional layer. Further, we aim to provide users the flexibility to interpret different levels of biomarkers through graph node pooling and several innovative loss terms to regulate the pooling operation. In addition, different from much of the GNN literature [47,34] where populational graphs based on fMRI are modeled by treating each subject as a node on the graph, we model each subject’s brain as one graph and each brain ROI as a node to learn ROI-based graph embeddings. Specifically, our framework jointly learns ROI clustering and the whole-brain fMRI prediction. This not only reduces preconceived errors, but also learns particular clustering patterns associated with the other quantitative brain image analysis tasks. Specifically, from estimated model parameters, we can retrieve ROI clustering patterns. Also, our GNN design facilitates model interpretability by regulating intermediate outputs with *a novel loss term for enforcing similarity of pooling scores*, which provides the flexibility to choose between individual-level and group-level explanations.

A preliminary version of this work, *Pooling Regularized Graph Neural Network (PR-GNN) for fMRI Biomarker Analysis* [39] was presented at the 22st International Conference on Medical Image Computing and Computer Assisted Intervention. This paper extends the preliminary version by designing novel graph convolutional layers and analyzing a new dataset and task.

## 2 BrainGNN

### 2.1 Notations

First we parcellate the brain into *N* ROIs based on its T1 structural MRI. We define ROIs as graph nodes 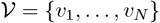 and the nodes are preordered. As brain ROIs can be aligned by brain parcellation atlases based on their locations in the structure space, we define the brain graphs as ordered aligned graphs. We define an undirected weighted graph as 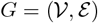, where 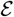 is the edge set, i.e., a collection of (*v_i_*, *v_j_*) linking vertices from *v_i_* to *v_j_*. In our setting, *G* has an associated node feature set and can be represented as matrix 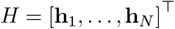, where **h**_i_ is the feature vector associated with node *v_i_*. For every edge connecting two nodes, 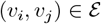, we have its strength 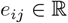 and *e_ij_* > 0. We also define *e_ij_* = 0 for 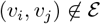 and therefore the adjacency matrix 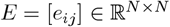 is well defined. We also list all the notations in Table 1.

**Table 1:**
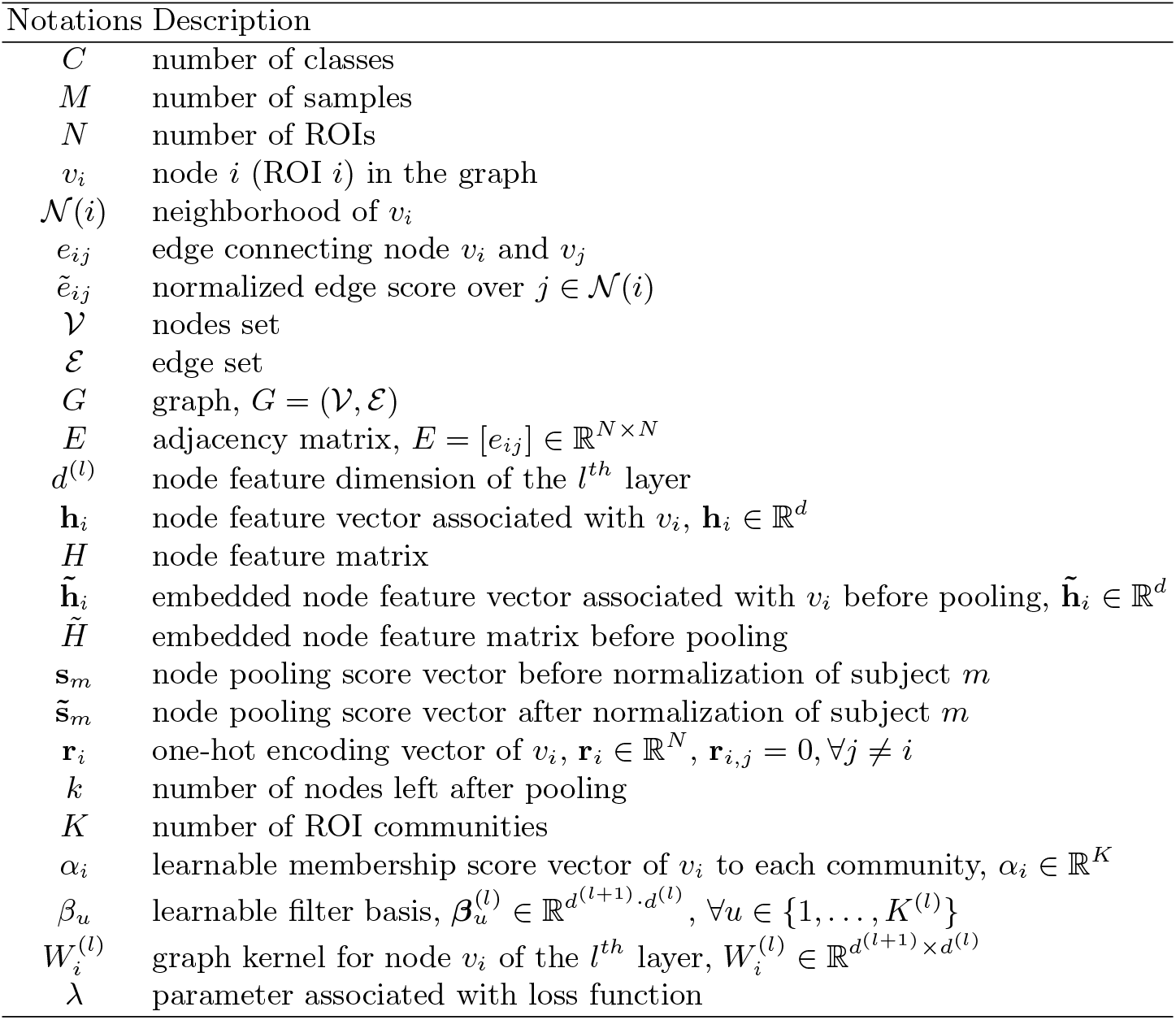
Notations used in the paper.

| Notations | Description |
| --- | --- |
| $C$ | number of classes |
| $M$ | number of samples |
| $N$ | number of ROIs |
| $v_i$ | node $i$ (ROI $i$) in the graph |
| $\mathcal{N}(i)$ | neighborhood of $v_i$ |
| $e_{ij}$ | edge connecting node $v_i$ and $v_j$ |
| $\tilde{e}_{ij}$ | normalized edge score over $j \in \mathcal{N}(i)$ |
| $\mathcal{V}$ | nodes set |
| $\mathcal{E}$ | edge set |
| $G$ | graph, $G = (\mathcal{V}, \mathcal{E})$ |
| $E$ | adjacency matrix, $E = [e_{ij}] \in \mathbb{R}^{N \times N}$ |
| $d^{(l)}$ | node feature dimension of the $l^{th}$ layer |
| $\mathbf{h}_i$ | node feature vector associated with $v_i$, $\mathbf{h}_i \in \mathbb{R}^d$ |
| $H$ | node feature matrix |
| $\tilde{\mathbf{h}}_i$ | embedded node feature vector associated with $v_i$ before pooling, $\tilde{\mathbf{h}}_i \in \mathbb{R}^d$ |
| $\tilde{H}$ | embedded node feature matrix before pooling |
| $\mathbf{s}_m$ | node pooling score vector before normalization of subject $m$ |
| $\tilde{\mathbf{s}}_m$ | node pooling score vector after normalization of subject $m$ |
| $\mathbf{r}_i$ | one-hot encoding vector of $v_i$, $\mathbf{r}_i \in \mathbb{R}^N$, $\mathbf{r}_{i,j} = 0, \forall j \neq i$ |
| $k$ | number of nodes left after pooling |
| $K$ | number of ROI communities |
| $\alpha_i$ | learnable membership score vector of $v_i$ to each community, $\alpha_i \in \mathbb{R}^K$ |
| $\beta_u$ | learnable filter basis, $\beta_u^{(l)} \in \mathbb{R}^{d^{(l+1)} \cdot d^{(l)}}$, $\forall u \in \{1, \dots, K^{(l)}\}$ |
| $W_i^{(l)}$ | graph kernel for node $v_i$ of the $l^{th}$ layer, $W_i^{(l)} \in \mathbb{R}^{d^{(l+1)} \times d^{(l)}}$ |
| $\lambda$ | parameter associated with loss function |

### 2.2 Architecture Overview

Classification on graphs is achieved by first embedding node features into a low-dimensional space, then coarsening or pooling nodes and summarizing them. The summarized vector is then fed into a multi-layer perceptron (MLP). We train the graph convolutional/pooling layers and the MLP in an end-to-end fashion. Our proposed network architecture is illustrated in Fig. (2a). It is formed by three different types of layers: graph convolutional layers, node pooling layers and a readout layer. Generally speaking, GNNs inductively learn a node representation

**Fig. 2:**
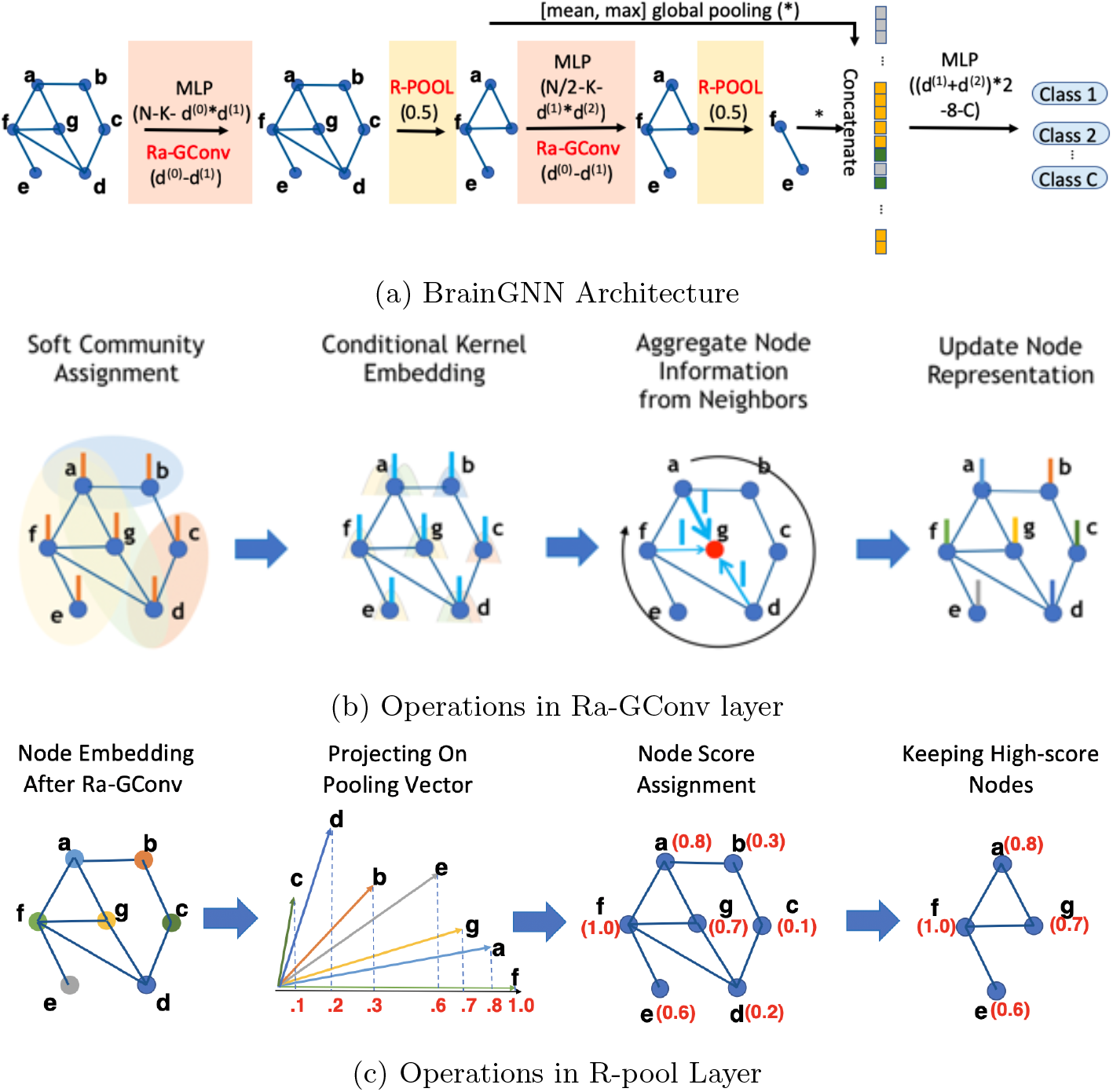
(a) introduces the BrainGNN architecture that we propose in this work. BrainGNN is composed of blocks of Ra-GConv layers and R-pool layers. It takes graphs as inputs and outputs graph-level predictions. (b) shows how the Ra-GConv layer embeds node features. First, nodes are softly assigned to communities based on their membership scores to the communities. Each community is associated with a different basis vector. Each node is embedded by the particular basis vectors based on the communities that it belongs to. Then, by aggregating a node’s own embedding and its neighbors’ embedding, the updated representation is assigned to each node on the graph. (c) shows how R-pool selects nodes to keep. First, all the nodes’ representations are projected to a learnable vector. The nodes with large projected values are retained with their corresponding connections.

by recursively transforming and aggregating the feature vectors of its neighboring nodes.

A **graph convolutional layer** is used to probe the graph structure by using edge features, which contain important information about graphs. For example, the weights of the edges in brain fMRI graphs can represent the relationship between different ROIs.

Following [51], we define 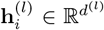 as the features for the *i^th^* node in the *l^th^* layer, where *d*^(*l*)^ is the dimension of the *l^th^* layer features. The propagation model for the forward-pass update of node representation is calculated as:

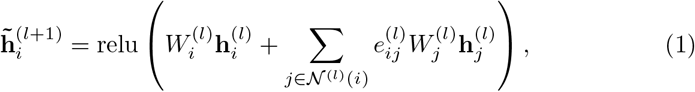

where 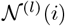 denotes the set of indices of neighboring nodes of node *v_i_*, 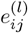 denotes the features associated with the edge from *v_i_*, to *v_j_*, 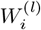 denote the model’s parameters to be learned. The first layer is operated on the original graph, i.e. 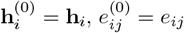. To avoid increasing the scale of output features, the edge features need to be normalized, as in GAT [55] and GNN [36]. Due to the aggregation mechanism, we normalize the weights by 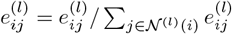.

A **node pooling** layer is used to reduce the size of the graph, either by grouping the nodes together or pruning the original graph *G* to a subgraph *G_s_* by keeping some important nodes only. We will focus on the pruning method, as it is more interpretable and can help detect biomarkers.

A **readout** layer is used to summarize the node feature vectors 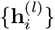 into a single vector **z**^(*l*)^ which is finally fed into a classifier for graph classification.

### 2.3 Layers in BrainGNN

In this section, we provide insights and highlight the innovative design aspects of our proposed BrainGNN architecture.

#### ROI-aware Graph Convolutional Layer

##### Overview

We propose an ROI-aware graph convolutional layer (Ra-GConv) with two insights. First, when computing the node embedding, we allow Ra-GConv to learn different embedding weights in graph convolutional kernels conditioned on the ROI (geometrically distributed information of the brain), instead of using the same weights *W* on all the nodes as shown in Eq. (1). In our design, the weights *W* can be decomposed as a linear combination of the basis set, where each basis function represents a community. Second, we include edge weights for message filtering, as the magnitude of edge weights presents the connection strength between two ROIs. We assume that more closely connected ROIs have a larger impact on each other.

##### Design

We begin by assuming the graphs have additional regional information and the nodes of the same region from different graphs have similar properties. We propose to encode the regional information into the embedding kernel function for the nodes. Given node *i*’s regional information **r**_*i*_, such as the node’s coordinates in a mesh graph, we propose to learn the vectorized embedding kernel 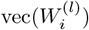 based on **r**_*i*_ for the *l^th^* Ra-GConv layer:

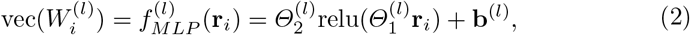

where the MLP with parameters 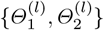 maps **r**_*i*_ to a *d*^(*l*+1)^ · *d*^(*l*)^ dimensional vector then reshapes the output to a *d*^(*l*+1)^ × *d*^(*l*)^ matrix 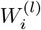 and **b**^(*l*)^ is the bias term in the MLP.

Given a brain parcellated into *N* ROIs, we order the ROIs in the same manner for all the brain graphs. Therefore, the nodes in the graphs of different subjects are aligned. However, the convolutional embedding should be independent of the ordering methods. Given an ROI ordering for all the graphs, we use one-hot encoding to represent the ROI’s location information, instead of using coordinates, because the nodes in the brain are aligned well. Specifically, for node *v_i_*, its ROI representation **r**_*i*_ is a *N*-dimensional vector with 1 in the *i^th^* entry and 0 for the other entries. Assume that 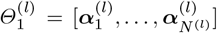, where *N*^(*l*)^ is the number of ROIs in the *l^th^* layer, 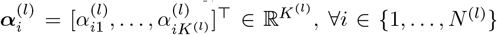, ∀*i* ∈{1,…, *N*^(*l*)^}, where *K*^(*l*)^ can be seen as the number of clustered communities for the *N*^(*l*)^ ROIs. Assume 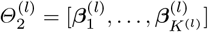 with 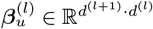, ∀*u* ∈{1,…, *K*^(*l*)^}. Then Eq. (2) can be rewritten as

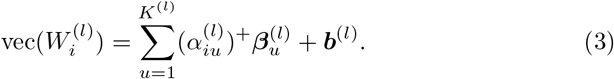

We can view 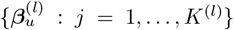 as a basis and 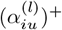 as the coordinates. From another perspective, 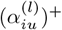 can be seen as the non-negative assignment score of ROI *i* to community *u*. If we train different embedding kernels for different ROIs for the *l^th^* layer, the total parameters to be learned will be *N*^(*l*)^*d*^(*l*)^*d*^(*l*+1)^. Usually we have *K*^(*l*)^ ≪ *N*^(*l*)^. By Eq. (3), we can reduce the number of learnable parameters to *K*^(*l*)^*d*^(*l*)^*d*^(*l*+1)^ + *N*^(*l*)^*K*^(*l*)^ parameters, while still assigning a separate embedding kernel for each ROI. The ROIs in the same community will be embedded by the similar kernel so that nodes in different communities are embedded in different ways.

As the graph convolution operations in [23], the node features will be multiplied by the edge weights, so that neighbors connected with stronger edges have a larger influence.

#### ROI-topK Pooling Layer

##### Overview

To perform graph-level classification, a layer for dimensionality reduction is needed since the number of nodes and the feature dimension per node are both large. Recent findings have shown that some ROIs are more indicative of predicting neurological disorders than the others [31,5], suggesting that they should be kept in the dimensionality reduction step. Therefore the node (ROI) pooling layer (R-pool) is designed to keep the most indicative ROIs while removing *noisy* nodes, thereby reducing the dimensionality of the entire graph.

##### Design

To make sure that down-sampling layers behave idiomatically with respect to different graph sizes and structures, we adopt the approach in [11] and [21] for reducing graph nodes. The choice of which nodes to drop is determined based on projecting the node features onto a learnable vector 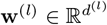. The nodes receiving lower scores will experience less feature retention. We denote 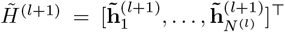, where *N*^(*l*)^ is the number of nodes at the *l^th^* layer. Fully written out, the operation of this pooling layer (computing a pooled graph, 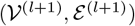, from an input graph, 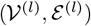), is expressed as follows:

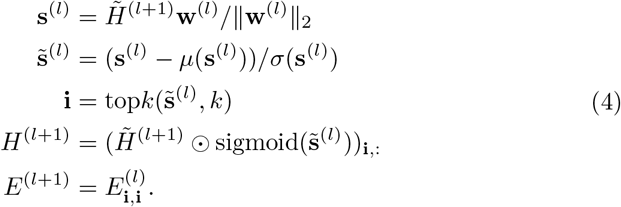

Here || · || is the *L*_2_ norm, *μ* and *σ* take the input vector and output the mean and standard deviation of its elements. The notation top*k* finds the indices corresponding to the largest *k* elements in score vector 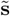. ⊙ is (broadcasted) element-wise multiplication, and (·)_**i,j**_ is an indexing operation which takes elements at row indices specified by **i** and column indices specified by **j** (colon denotes all indices). The pooling operation retains sparsity by requiring only a projection, a point-wise multiplication and a slicing into the original features and adjacency matrix. Different from [11], we added element-wise score normalization 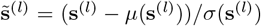, which is important for calculating the loss functions in Section 2.4.

###### Readout Layer

Lastly, we seek a “flattening” operation to preserve information about the input graph in a fixed-size representation. Concretely, to summarize the output graph of the *l^th^* conv-pool block, 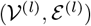, we use

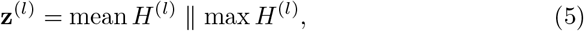

where 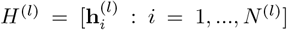, mean and max operate element-wisely, and || denotes concatenation. To retain information of a graph in a vector, we concatenate both mean and max summarization for a more informative graph-level representation. The final summary vector is obtained as the concatenation of all those summaries (i.e. **z** = **z**^(1)^ || **z**^(2)^ || ··· || **z**^(*L*)^) and it is submitted to a MLP for obtaining final predictions.

###### Putting Layers Together

All in all, the architecture (as shown in Fig. 2a) consists of two kinds of layers — Ra-GConv layers shown in the pink blocks and R-pool layer shown in the yellow blocks. The input is a weighted graph with its node attributes constructed from fMRI. We form a two-layer GNN block starting with ROI-aware node embedding by the proposed Ra-GConv layer in Section 2.3, followed by the proposed R-pool layer in Section 2.3. The whole network sequentially concatenates these GNN blocks, and readout layers are added after each GNN block. The final summary vector concatenates all the summaries from the readout layers, and an MLP is applied after that to give final predictions.

### 2.4 Loss Functions

The classification loss is the cross entropy loss:

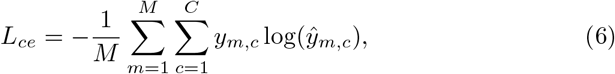

where *M* is the number of instances, *C* is the number of classes, *y_mc_* is the ground truth label and 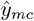 is the model output.

Now we describe the loss terms designed to regulate the learning process and control the interpretability.

#### Unit loss

As we mentioned in Section 2.3, we project the node representation to a learnable vector 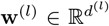. The learnable vector **w**^(*l*)^ can be arbitrarily scaled while the pooling scores 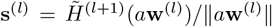 remain the same with non-zero scalar 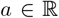. This suggests an identifiability issue, i.e. multiple parameters generate the same distribution of the observed data. To remove this issue, we add a constraint that **w**^(*l*)^ is a unit vector. To avoid the problem of identifiability, we propose unit loss:

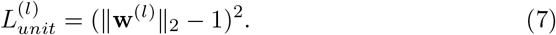

#### Group-level consistency loss

We propose group-level consistency (GLC) loss to force BrainGNN to select similar ROIs in a R-pool layer for different input instances. This is because for some applications, users may want to find the common patterns/biomarkers for a certain neuro-prediction task. Note that 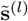 in Eq. (4) is computed from the input *H*^(*l*)^ and they change as the layer goes deeper for different instances. Therefore, for different inputs *H*^(*l*)^, the selected entries of 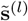 may not correspond to the same set of nodes in the original graph, so it is not meaningful to enforce similarity of these entries. Thus, we only use the GLC loss regularization for 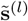 vectors after the first pooling layer.

Now, we mathematically describe the novel GLC loss. In each training batch, suppose there are *M* instances, which can be partitioned into *C* subsets based on the class labels, 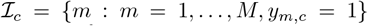, for *c* = 1,…, *C*. And *y_m,c_* = 1 indicates the *m^th^* instance belongs to class *c*. We form the scoring matrix for the instances belonging to class *c* as 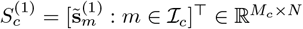, where 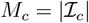. The GLC loss can be expressed as:

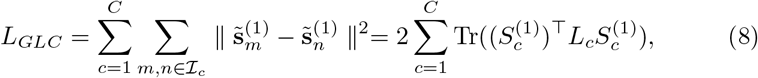

where *L_c_* = *D_c_*–*W_c_* is a symmetric positive semidefinite matrix, *W_c_* is a *M_c_* × *M_c_* matrix with values of 1, *D_c_* is a *M_c_* × *M_c_* diagonal matrix with *M_c_* as diagonal elements [57], *m* and *n* are the indices for instances. Thus, Eq. (8) can be viewed as calculating pairwise pooling score similarities of the instances.

#### TopK pooling loss

We propose TopK pooling (TPK) loss to encourage reasonable node selection in R-pool layers. In other words, we hope the top *k* selected indicative ROIs should have significantly different scores than those of the unselected nodes. Ideally, the scores for the selected nodes should be close to 1 and the scores for the unselected nodes should be close to 0. To achieve this, we rank 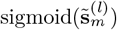 for the *m*th instance in a descending order, denote it as 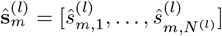, and apply a constraint to all the *M* training instances to make the values of 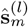 more dispersed. In practice, we define TPK loss using binary cross-entropy as:

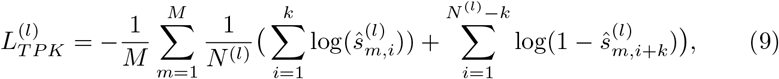

We show the kernel density estimate plots of normalized node pooling scores (indication of the importance of the nodes) changing over the training epoch in Fig. 3 when 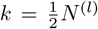. It is clear to see that the pooling scores are more dispersed over time, Hence the top 50% selected nodes have significantly higher importance scores than the unselected ones. In the experiments below, we further demonstrate the effectiveness of this loss term in an ablation study. For now, we finalize our loss function below.

**Fig. 3:**
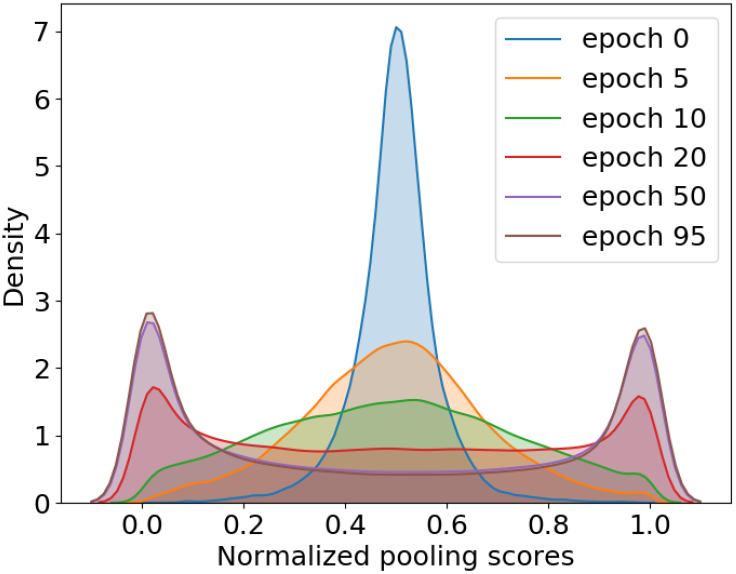
The change of the distribution of node pooling scores 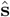 of the 1st R-pool layer over 100 training epochs presented using kernel density estimate plots. With TopK pooling (TPK) loss, the node pooling scores of the selected nodes and those of the unselected nodes become significantly separate.

Finally, the final loss function is formed as:

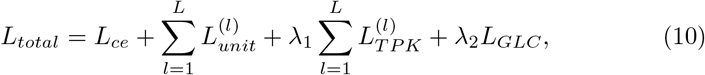

where λ’s are tunable hyper-parameters, *l* indicates the *l^th^* GNN block and *L* is the total number of GNN blocks. To maintain a concise loss function, we do not have tunable hyper-parameters for *L_ce_* and 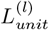. We observed that the unit loss 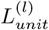 can quickly decrease to a small number close to zero. Empirically, this term and the cross entropy term *L_ce_* already have the same magnitude (suppose the latter ranges from – log(0.5) to – log(1)). If the unit loss is much larger than the cross entropy term, the entire loss function will penalize it more and force it to have the same magnitude as the cross entropy. Also, since **w**^(*l*)^ can be arbitrarily scaled without changing the output, the optimization can scale it to reduce the entire loss without affecting the other terms.

### 2.5 Interpretation from BrainGNN

#### Community Detection from Convolutional Layers

The important contribution of our proposed ROI-aware convolutional layer is the implied community clustering patterns in the graph. Discovering brain community patterns is critical to understanding co-activation and interaction in the brain. Revisiting Eq. (3) and following [40], 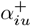 provides the membership of ROI *i* to community *u*. The community assignment is soft and overlaid. Specifically, we consider region *i* belongs to community *u* if 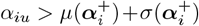. This gives us a collection of community indices indicating region membership {***i**_u_* ⊂ {1,…, *N*}: *u* = 1,…,*K*}.

#### Biomarker Detection from Pooling Layers

Without the added TPK loss (Eq. (9)), the significance of the nodes left after pooling cannot be guaranteed. With TPK loss, pooling scores are more dispersed over time, hence the selected nodes have significantly higher importance scores than the unselected ones.

The strength of the GLC loss controls the trade-off between individual-level interpretation and group-level interpretation. On the one hand, for precision medicine, individual-level biomarkers are desired for planning targeted treatment. On the other hand, group-level biomarkers are essential for understanding the common characteristic patterns associated with the disease. We can tune the coefficient λ_2_ to control different levels of interpretation. Large λ_2_ encourages selecting similar nodes, while small λ_2_ allows various node selection results for different instances.

## 3 Experiments and Results

### 3.1 Datasets

Two independent datasets are used: the Biopoint Autism Study Dataset (Bio-point) [56] and the Human Connectome Project (HCP) 900 Subject Release [54]. For the Biopoint dataset, the aim is to classify Autism Spectrum Disorder (ASD) and Healthy Control (HC). For the HCP dataset, like the recent work [59,61,43], the aim is to decode and map cognitive states of the human brain. Thus, we classify 7 task states - gambling, language, motor, relational, social, working memory (WM), and emotion, then infer the decoded task-related salient ROIs from interpretation. The HCP states classification task helps validate our interpretation results (will discuss in Section 3.5). These represent two key examples of task-based paradigms that will illustrate the power and portability of our approach.

#### Biopoint Dataset

The Biopoint Autism Study Dataset [56] contains task fMRI scans for ASD and neurotypical healthy controls (HCs). The subjects perform the “biopoint” task, viewing point-light animations of coherent and scrambled biological motion in a block design [31] (24*s* per block). The fMRI data are preprocessed using the pipeline described in [56], and includes the removal of subjects with significant head motion during scanning. This results in 72 ASD children and 43 age-matched (*p* > 0.124) and IQ-matched (*p* > 0.122) neurotypical HCs. We insured that the head motion parameters are not significantly different between the groups. There are more male subjects than female samples, similar to the level of ASD prevalence in the population [18,28]. The first few frames are discarded, resulting in 146 frames for each fMRI sequence.

The Desikan-Killiany [14] atlas is used to parcellate brain images into 84 ROIs. The mean time series for each node is extracted from a random 1/3 of voxels in the ROI (given an atlas) by bootstrapping. We use Pearson correlation coefficient as node features (i.e a vector of Pearson correlation coefficients to all ROIs). Edges are defined by thresholding (in practice, we use top 10% positive which guarantees no isolated nodes in the graph) partial correlations to achieve sparse connections. We use partial correlation to build edges for the following two reasons: 1) due to the over-smoothing effect of the general graph neural networks for densely connected graphs [46,10], it is better to avoid dense graphs and partial correlation tends to lead to sparse graphs; 2) Pearson correlation and partial correlation are different measures of fMRI connectivity; we aggregate them by using one to build edge connections and the other to build node features. This is motivated by recent multi-graph fusion works for neuroimaging analysis that aim to capture different brain activity patterns by leveraging different correlation matrices [63,20]. Hence, node features are 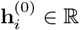. Each fMRI dataset is augmented 30 times by spatially resampling the fMRI bold signals [16]. Specifically, we randomly sample 1/3 of the voxels within an ROI to calculate the mean time series. This sampling process is repeated 30 times, resulting in 30 graphs for each fMRI image instance.

#### HCP Dataset

For this dataset, we restrict our analyses to those individuals who participated with full length of scan, whose mean frame-to-frame displacement is less than 0.1 mm and whose maximum frame-to-frame displacement is less than 0.15 mm (n=506; 237 males; ages 22–37). This conservative threshold for exclusion due to motion is used to mitigate the substantial effects of motion on functional connectivity.

We process the HCP fMRI data with standard methods (see [17] for more details) and parcellated into 268 nodes using a whole-brain, functional atlas defined in a separate sample (see [25] for more details). For the easy of validating the task-related function key words, our classification focuses on task fMRI in the HCP dataset. Task functional connectivity is calculated based on the raw task time series: the mean time series of each node pair were used to calculate the Pearson correlation and partial correlation. We define a weighted undirected graph with 268 nodes per individual per task condition resulting in 3542 = 506 × 7 graphs in total. The same graph construction method as for the Biopoint data is used. Hence, node feature 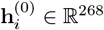.

### 3.2 Experimental Setup

We trained and tested the algorithm on Pytorch in the Python environment using a NVIDIA Geforce GTX 1080Ti with 11GB GPU memory. The model architecture was implemented with 2 conv layers and 2 pooling layers as shown in Fig. (2a), with parameter *N* =84, *K*^(0)^ = *K*^(1)^ = 8, *d*^(0)^ = 84, *d*^(1)^ = 16, *d*^(2)^ = 16, *C* = 2 for the Biopoint dataset and *N* = 268, *K*^(0)^ = *K*^(1)^ = 8, *d*^(0)^ = 268, *d*^(1)^ = 32, *d*^(2)^ = 32, *C* = 7 for HCP dataset. In our work, we set *k* in Eq 4 as half of nodes in that layer, namely the dropout rate is 0.5. The motivation of *K* = 8 comes from the eight functional networks defined by Finn et al. [17], because these 8 networks show key brain functionality relevant to our tasks.

We will discuss the variation of λ_1_ and λ_2_ in Section 3.3. We first hold 1/5 data as the testing set and then randomly split the rest of the dataset into a training set (3/5 data), and a validation set (1/5 data) used to determine the hyperparameters. The graphs from a single subject can only appear in either the training, validation or testing set. Specifically, for the Biopoint dataset, each training set contains 2070 graphs (69 subjects and 30 graphs per subject), each validation set contains 690 graphs (23 subjects and 30 graphs per subject), and the testing set contains 690 graphs (23 subjects, and 30 graphs per subject). For the HCP dataset, each training set contains 2121 or 2128 graphs (303 or 304 subjects, and 7 graphs per subject), each validation set contains 707 or 714 graphs (101 or 102 subjects and 714 graphs per subject), and the testing set contains 690 graphs (102 subjects and 7 graphs per subject). In this section, we use training and validation sets only to study λ_1_ and λ_2_. Adam was used as the optimizer. We trained BrainGNN for 100 iterations with an initial learning rate of 0.001 and annealed to half every 20 epochs. Each batch contained 400 graphs for Biopoint data and 200 graphs for HCP data. The weight decay parameter was 0.005.

### 3.3 Hyperparameter Discussion and Ablation Study

#### Hyperparameter discussion setup

To check how the hyperparameters affect the performance, we tune λ_1_ and λ_2_ in the loss function using the training and validation sets. Recalling our intuition of designing TPK loss and GLC loss described in Section 2.4, large λ_1_ (TPK loss) encourages more separable node importance scores for selected and unselected nodes after pooling, and λ_2_ (GLC loss) controls the similarity of the nodes selected by different instances (hence controls the level of interpretability between individual-level and group-level). Small λ_2_ would result in variant individual-specific patterns, while large λ_2_ would force the model to learn common group-level patterns. As task classification on HCP could achieve consistently high accuracy over the parameter variations, we only show the results on the Biopoint validation sets generated from five random splits in Fig. 4.

**Fig. 4:**
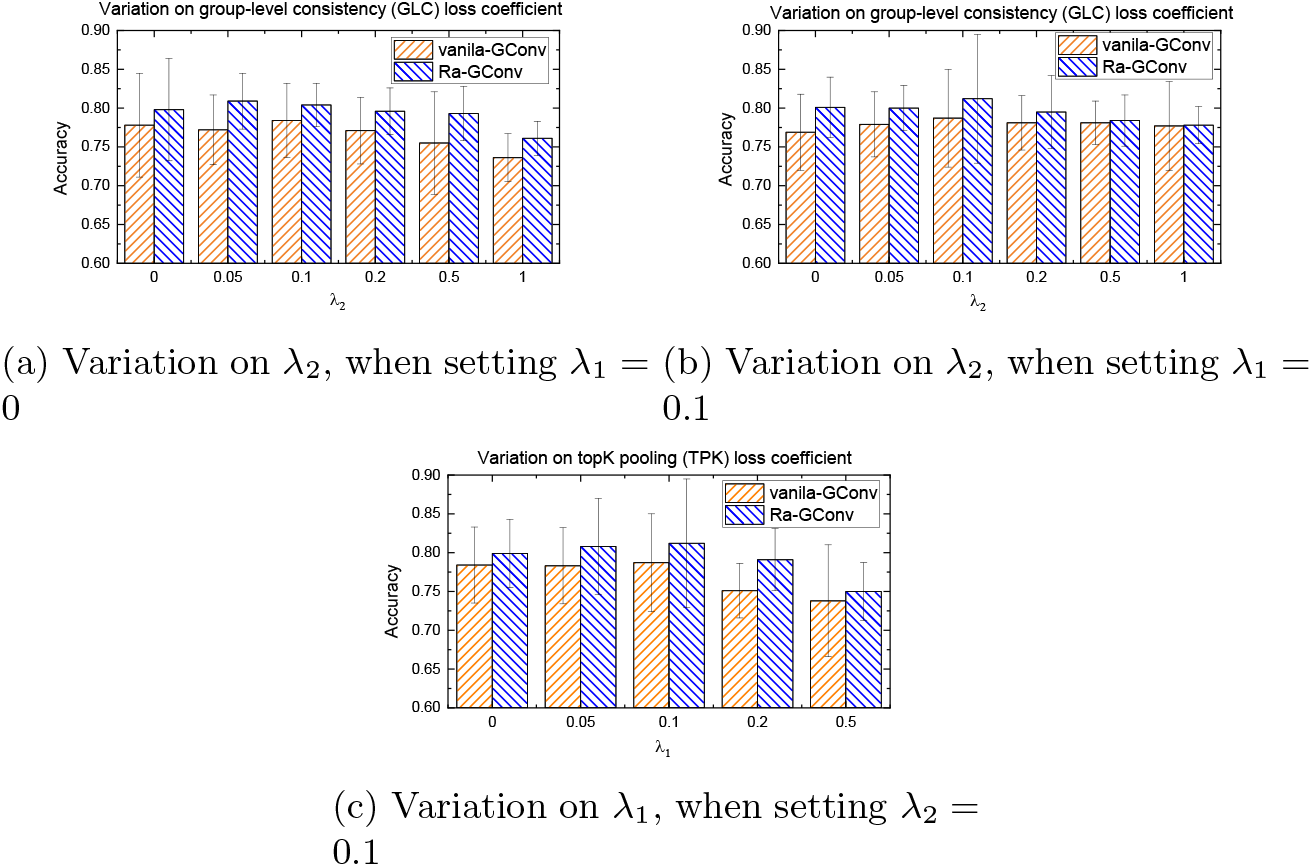
Comparison of Ra-GConv with vanilla-GConv and effect of coefficients of total loss in terms of accuracies on the validation sets.

#### Ablation study setup

To investigate the potential benefits of our proposed ROI-aware graph convolutional mechanism, we perform ablation studies. Specifically, we compare our proposed Ra-GConv layer with the strategy of directly learning embedding kernels *W* (without ROI-aware setting), which is denoted as ‘vanilla-GConv’.

#### Results

We evaluate the best classification accuracy on the validation sets in the 5-fold cross-validation setting. Due to the expensive cost involved in training deep learning models, we adopt an empirical way that first tunes λ_2_ with λ_1_ fixed to 0 or 0.1 and then tunes λ_1_ given the determined λ_2_.

First, we investigate the effects of λ_2_ on the accuracy with λ_1_ fixed to 0. The results are shown in Fig. 4a. We notice that the results are stable to the variation of λ_2_ in the range 0–0.5. When λ_2_ = 1, the accuracy drops. The accuracy reaches the peak when λ_2_ = 0.1. As the other deep learning models behave, BrainGNN is overparameterized. Without regularization (λ_2_ = 0), the model is easier to overfit to the training set, while large regularization of GLC might result in underfitting to the training set.

Second, we fix λ_1_ = 0.1 and varied λ_2_ again. As the results presented in Fig. 4b show, the accuracy drops if we increase λ_2_ after 0.2, which follows the same trend in Fig. 4a. However, the accuracy under the setting of λ_2_ = 0 is better than that in Fig. 4a. This is probably because the λ_1_ terms can work as regularization and mitigate the overfitting issue.

Last, we fix λ_2_ = 0.1 and vary λ_1_ from 0 to 0.5. As the results in Fig. 4c show, when we increased λ_1_ to 0.2 and 0.5, the accuracy slightly dropped.

For ablation study, as the results in Fig. 4 show, we can conclude that Ra-GConv overall outperformed the vanilla-GConv strategy under all the parameter settings. The reason could be better node embedding from multiple embedding kernels in the Ra-GConv layers, as the vanilla-GConv strategy treats ROIs (nodes) identically and used the same kernel for all the ROIs. Hence, we claim that Ra-GConv can better characterize the heterogeneous representations of brain ROIs.

Based on the results of tuning λ_1_ and λ_2_ on the validation sets, we choose the best setting of λ_1_ = λ_2_ = 0.1 for the following baseline comparison experiments. We report the results on the held-out testing set.

### 3.4 Comparison with Baseline Methods

We compare our method with traditional machine learning (ML) methods and state-of-the-art deep learning (DL) methods to evaluate the classification accuracy. The ML baseline methods take vectorized correlation matrices as inputs, with dimension *N*^2^, where *N* is the number of parcellated ROIs. These methods included Random Forest (1000 trees), SVM (RBF kernel), and MLP (2 layers with 20 hidden nodes). A variety of DL methods have been applied to brain connectome data, e.g. long short-term memory (LSTM) recurrent neural network [13], and 2D CNN [33,30], but they are not designed for brain graph analysis. Here we choose to compare our method with BrainNetCNN [33], which is designed for fMRI network analysis. We also compare our method with other GNN methods: GAT [55], GraphSAGE [26], and our preliminary version PR-GNN [39]. It is worth noting that GraphSAGE does not take edge weights in the aggregation step of the graph convolutional operation. The inputs of BrainNetCNN are correlation matrices. We follow the parameter settings indicated in the original paper [33]. The inputs and the settings of hidden layer nodes for the graph convolution, pooling and MLP layers of the alternative GNN methods are the same as BrainGNN. We also show the number of trainable parameters required by each method. We repeat the experiment and randomly split independent training, validation, and testing sets five times. Hyperparameters for baseline methods are also tuned on the validation sets and we report the results on the five testing sets in Table2.

**Table 2:** Comparison of the classification performance with different baseline machine learning models and state-of-the-art deep learning models.

|  |  | SVM | Random Forest | MLP | BrainNetCNN | GAT | GraphSAGE | PR-GNN | BrainGNN |
| --- | --- | --- | --- | --- | --- | --- | --- | --- | --- |
| Biopoint | Accuracy (%) | 62.80(4.92)<sup>a</sup> | 68.60(3.58) | 58.80(1.79) | 75.20(3.49) | 77.40(3.51) | 78.60(5.90) | 77.10(8.71) | **79.80(3.63)<sup>c</sup>** |
|  | F1 (%) | 60.08(3.91) | 63.97(4.95) | 55.25(9.49) | 65.58(14.48) | 75.08(5.19) | 75.55(7.03) | 75.20(7.01) | **75.80(6.03)** |
|  | Recall (%) | 60.20(4.49) | 71.11(8.12) | 61.00(4.85) | 66.20(10.85) | 71.60(6.07) | 75.20(6.46) | 78.26(10.28) | **72.60(5.64)** |
|  | Precision (%) | 60.00(3.81) | 67.80(5.36) | 53.40(12.52) | 65.60(17.95) | 79.40(8.02) | 76.20(8.11) | 76.50(14.32) | **79.60(8.59)** |
|  | Parameter (k) | 3 | 3 | 138 | 1438 | 16 | 6 | 6 | 41 |
| HCP | Accuracy (%) | 90.00(8.20) | 90.20(4.15) | 67.20(34.40) | 90.60(4.04) | 78.60(10.45) | 89.80(12.51) | 91.20(8.28) | **94.40(4.04)\*<sup>d</sup>** |
|  | F1 (%) | 90.20(5.81) | 90.14(5.55) | 63.49(41.80) | 90.96(3.50) | 77.00(11.58) | 88.60(13.19) | 91.09(8.35) | **94.34(3.27)\*** |
|  | Recall (%) | 89.57(8.04) | 90.06(7.35) | 67.97(41.66) | 91.12(4.13) | 78.60(10.45) | 89.43(12.43) | 91.00(8.95) | **94.29(3.73)\*** |
|  | Precision (%) | 90.85(9.35) | 90.22(4.77) | 62.97(42.47) | 90.81(3.27) | 91.20(3.32) | 87.80(14.02) | 91.14(8.52) | **94.40(3.59)\*** |
|  | Parameter (k) | 36 | 36 | 713 | 4547 | 34 | 12 | 12 | 96 |

<sup>a</sup> Classification accuracy, f1-score, recall and precision of the testing sets are reported in mean (standard deviation) format.
<sup>b</sup> The number of trainable parameters of each model is denoted.
<sup>c</sup> We boldfaced the results generated from our proposed BrainGNN.
<sup>d</sup> \* indicates significantly outperforming ( $p < 0.001$ under one tail two-sample t-test) all the alternative methods.

As shown in Table2, we report the comparison results using four different evaluation metrics, including accuracy, F1-score, recall and precision. We report the mean and standard deviation of the metrics on the five testing sets. We use validation sets to select the early stop epochs for the deep learning methods. On the HCP dataset, the performance of our BrainGNN significantly exceeds that of the alternative methods (*p* < 0.001 under one tail two-sample t-test). On the Biopoint dataset, as data augmentation are performed on all the data points for the consistency of cross validation and to improve prediction performance, we report the subject-wise metric through majority-voting on the predicted label from the augmented inputs. BrainGNN is significantly better than most of the alternative methods (*p* < 0.05 under one tail two-sample t-test) except for the previous version of our own work, PR-GNN and BrainGNN, although the mean values of all the metrics are consistently better than PR-GNN and BrainNetCNN. The improvement may result from two causes. First, due to the intrinsic complexity of fMRI, complex models with more parameters are desired, which also explains why CNN and GNN-based methods were better than SVM and random forest. Second, our model utilized the properties of fMRI and community structure in the brain network and thus potentially modeled the local integration more effectively. Compared to alternative machine learning models, BrainGNN achieved significantly better classification results on two independent task-fMRI datasets. Moreover, BrainGNN does not have the burden of feature selection, which is needed in traditional machine learning methods. Compared with MLP and CNN-based methods, GNN-based methods require less trainable parameters. Specifically, BrainGNN needs only 10 – 30% of the parameters of MLP and less than 3% of the parameters of BrainNetCNN. Our method requires less parameters and achieves higher data utility, hence it is more suitable as a deep learning tool for fMRI analysis, when the sample size is limited.

### 3.5 Interpretability of BrainGNN

A compelling advantage of BrainGNN is its ***built-in*** interpretability: (1) on the one hand, users can interpret salient brain regions that are informative to the prediction task at different levels; (2) on the other hand, BrainGNN clusters brain regions into prediction-related communities. We demonstrate (1) in Section 3.5-3.5 and (2) in Section 3.5. We show how our method can provide insights on the salient ROIs, which can be treated as disease-related biomarkers or fingerprints of cognitive states.

#### Individual- or Group-Level Biomarker

It is essential for a pipeline to be able to discover personal biomarkers and group-level biomarkers in different application scenarios, i.e. precision medicine and disease understanding. In this section, we discuss how to adjust λ_2_, the parameter associated with GLC loss, to manipulate the level of biomarker interpretation through training.

Our proposed R-pool can prune the uninformative nodes and their connections from the brain graph based on the learning tasks. In other words, only the salient nodes are kept/selected. We investigate how to control the similarity between the selected ROIs of different individuals by tuning λ_2_. As we discuss in Section 2.5, large λ_2_ encourages group-level interpretation (similar biomarkers across subjects) and small λ_2_ encourages individual-level interpretation (various biomarkers across subjects). But when λ_2_ is too large, the regularization might hurt the model accuracy (shown in Fig. 4). We put forth the hypothesis that meaningful interpretation is more likely to be derived from a model with high classification accuracy, as suggested in [27,2]. Intuitively, interpretation is trying to understand how a model makes a right decision rather than a wrong one when learning from a good teacher. We take the model with the highest accuracy for the interpretation experiment. Hence, the interpretation is restricted to models with fixed λ_1_ = 0.1 and varying λ_2_ from 0 to 0.5 according to our experiments in Section 3.3. Without losing the generalizability, we show the salient ROI detection results of 3 randomly selected ASD instances from the Biopoint dataset in Fig. 5. We show the remaining 21 ROIs after the 2nd R-pool layer (with pooling ratio = 0.5, 25% nodes left) and corresponding pooling scores. As shown in Fig. 5(a), when λ_2_ = 0, “overlapped areas” (defined as spatial areas where saliency values agree) among the three instances are rarely to be found. The various salient brain ROIs are biomarkers specific to each individual. Many clinical applications, such as personalized treatment outcome prediction or disease subtype detection, require learning the individual-level biomarkers to achieve the best predictive performance [8,6]. However, in some other applications, such as understanding the general pattern or mechanism associated with a cognitive task or disease, group-level biomarkers which highlight consistent explanations across individuals are important [3,56,50]. We can increase λ_2_ to achieve such group-level explanations. In Fig. 5(b-c), we circle the big “overlapped areas” across the three instances. By visually examining the salient ROIs, we find three “overlapped areas” in Fig. 5(b) and five “overlapped areas” in Fig. 5(c).

**Fig. 5:**
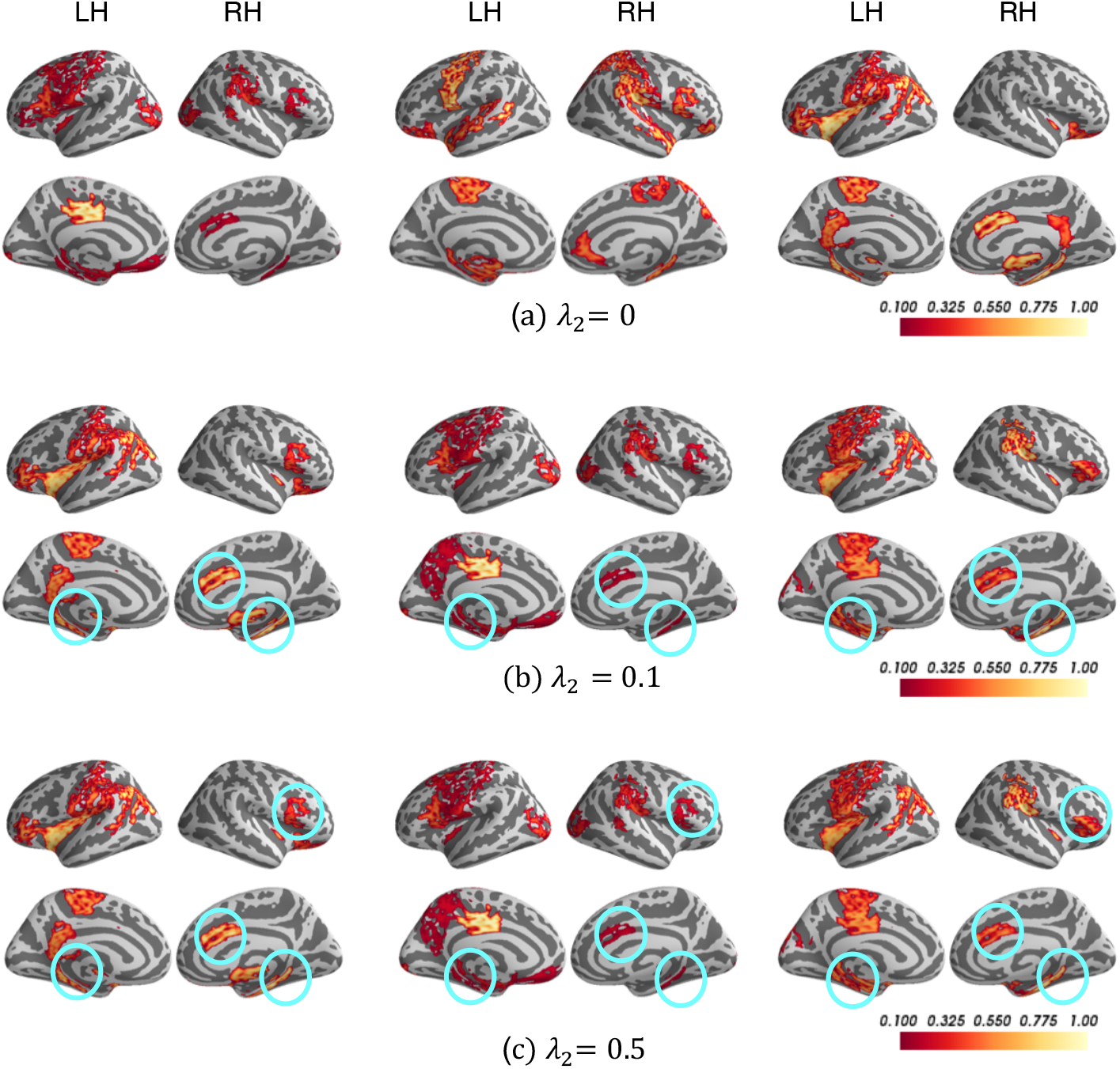
Interpretation results of Biopoint task. The selected salient ROIs of three different ASD individuals with different weights λ_2_ associated with group-level consistency term *L_GLC_*. The color bar ranges from 0.1 to 1. The bright-yellow color indicates a high score, while dark-red color indicates a low score. The commonly detected salient ROIs across different individuals are circled in blue.

#### Validating Salient ROIs

To demonstrate the effectiveness of the interpreted salient ROIs, we compare the biomarkers with existing literature studies. We average the node pooling scores after the 1st R-pool layer for all subjects per class and select the top salient ROIs as biomarkers for that class.

In Fig. 6, we display the salient ROIs (the top 21 ROIs, 21 = 84 × 0.5 × 0.5, where 84 is the total number of ROIs, and 0.5 is the pooling ratio of two R-pool layers) associated with HC and ASD separately. Putamen, thalamus, temporal gyrus and insular, occipital lobe are selected for HC; frontal gyrus, temporal lobe, cingulate gyrus, occipital pole, and angular gyrus are selected for ASD. Hippocampus and temporal pole are important for both groups. We name the selected ROIs as the biomarkers for identifying each group.

**Fig. 6:**
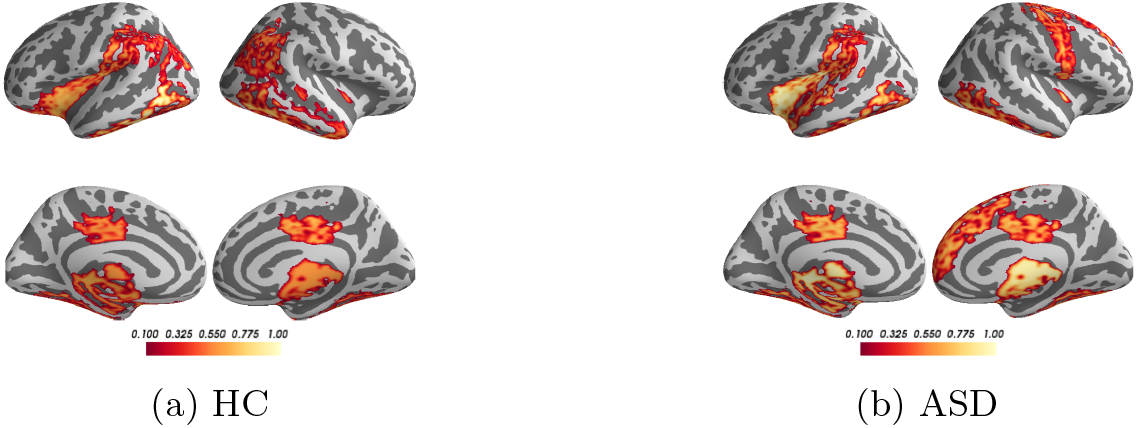
Interpretation results of Biopoint task. Interpreting salient ROIs (importance scores are denoted in colorbar) for classifying HC vs. ASD using BrainGNN.

The biomarkers for HC corresponded to the areas of clear deficit in ASD, such as social communication, perception, and execution. In contrast, the biomarkers for ASD map to implicated activation-exhibited areas in ASD: default mode network [9] and memory [7]. This conclusion is consistent both with behavioral observations when administering the fMRI paradigm and with a prevailing theory that ASD includes areas of cognitive strengths amidst the social deficits [48,53,29].

In Fig. 7(a-g), we list the salient ROIs associated with the seven tasks for the HCP dataset. To validate the neurological significance of the result, we used Neurosynth [64], a platform for fMRI data analysis. Neurosynth collects thousands of neuroscience publications and provides meta-analysis that gives keywords and their associated statistical images. The decoding function on the platform calculates the correlation between the input image and each functional keyword’s meta-analysis images. A high correlation indicates large association between the salient ROIs and the functional keywords. We selected the names of the tasks —‘gambling’, ‘language’, ‘motor’, ‘relational’, ‘social’, ‘working memory’ (WM) and ‘emotion’, as the functional keywords to be decoded. The heatmap in Fig. 8 illustrates the meta-analysis on functional keywords implied by the top salient regions corresponding to the seven tasks using Neurosynth. We define a state set, which is the same as the functional keywords set, as 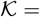 {‘gambling’,‘language’, ‘motor’, ‘relational’, ‘social’, ‘WM’, ‘emotion’}. In practice, given the interpreted salient ROIs associated with a functional state 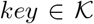, we generate the corresponding binary ROI mask. The mask is used as the input for Neurosynth analysis, which generates a vector of association scores between salient ROIs of *key* and all the keywords in 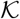 as shown in each row of Fig. 8. To facilitate visualization, we divide each value by the maximum absolute value of each column for normalization. If the diagonal value (from bottom left to top right) is 1, it indicates the interpreted salient ROIs reflect its real task state. The finding in Fig. 8 suggests that our algorithm can identify ROIs that are key to distinguish between the 7 tasks. For example, the anterior temporal lobe and temporal parietal regions, which are selected for the social task, are typically associated with social cognition in the literature [42,49].

**Fig. 7:**
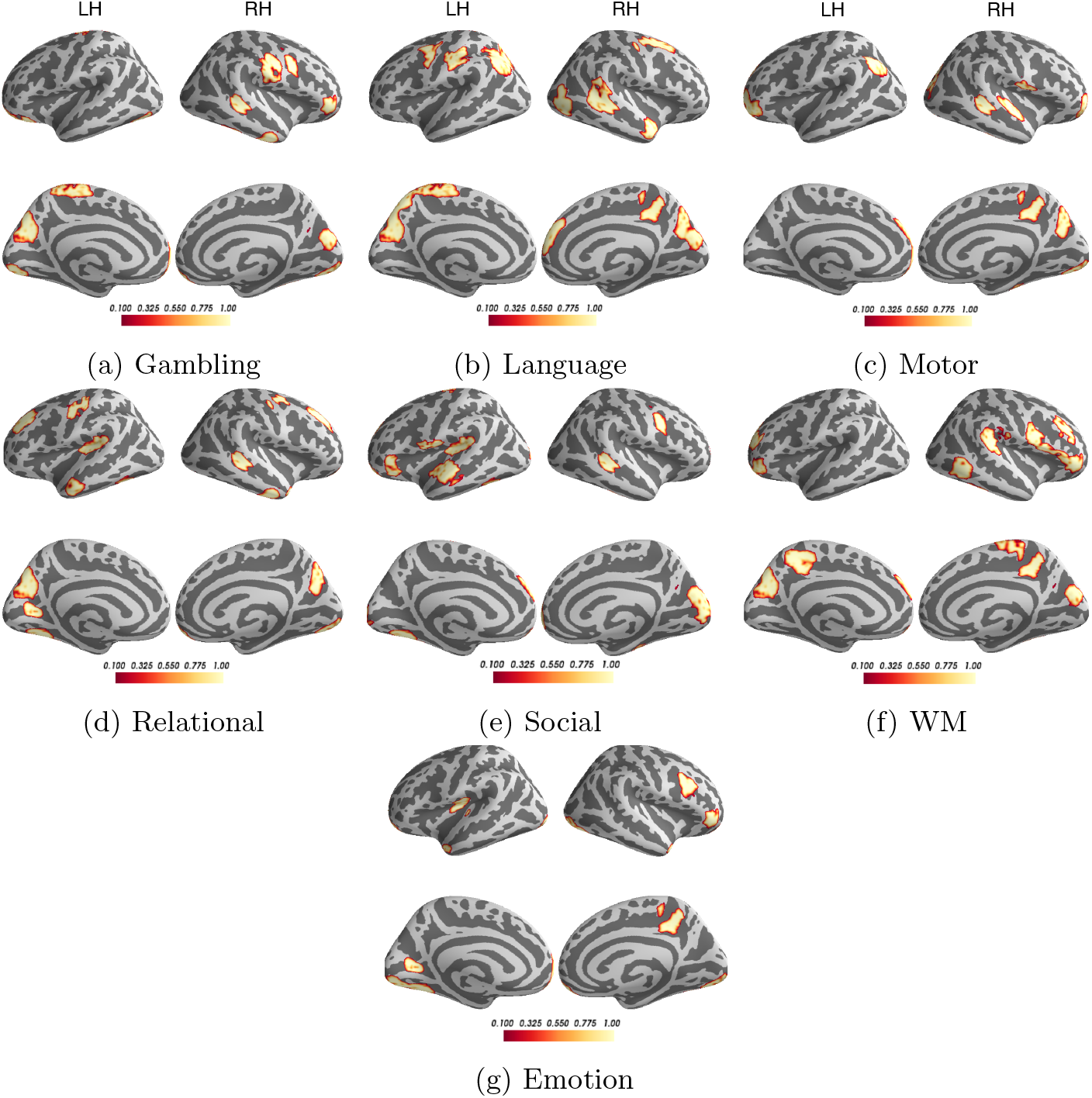
Interpretation results of HCP task. Interpreting salient ROIs (importance scores are denoted in color-bar) associated with classifying seven tasks.

**Fig. 8:**
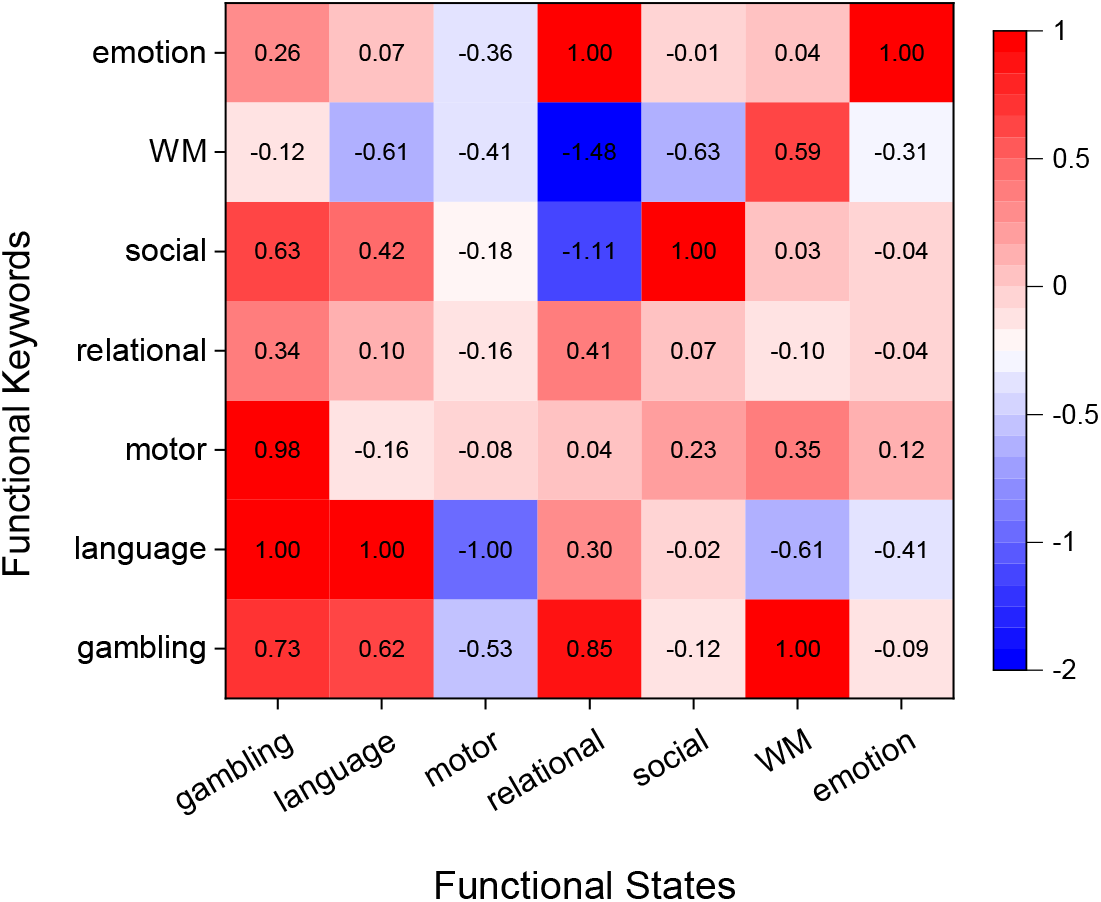
The correlation coefficient decoded by NeuroSynth (normalized by dividing it by the largest absolute value of *each column* for better visualization) between the interpreted biomarkers and the functional keywords for each functional state. A large correlation (in red) along *each column* indicates large association between the salient ROIs and the functional keyword. Large values (in red) on the diagonal from left-bottom to right-top indicate reasonable decoding; especially a value of 1.00 on the diagonal means that the interpreted salient ROIs of the task state are most correlated with the keywords of that state among all possible states in Neurosynth.

#### Node Clustering Patterns in Ra-GConv layer

From the best fold of each dataset, we cluster all the ROIs based on the kernel parameter 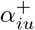 (learned in Eq. (3)) of the 1st Ra-GConv layer, which indicates the membership score of region *i* for community *u*. We show the node clustering results for the Biopoint and HCP data in Fig. 9a and Fig. 9b respectively. For the clustering results on the ASD classification task (shown in Fig. 9a), we observed the spatial aggregation patterns of each community, while the community clustering results on HCP task (shown in Fig. 9b) do not form similar spatial patterns. The different community clustering results reveal that the brain ROI community patterns are likely different depending on the tasks. Fig. 10 shows that the membership scores (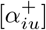 matrices) are not uniformly distributed across the communities and only one or a few communities have significantly larger scores than the other communities for a given ROI. This corroborates the necessity of using different kernels to learn node representation by forming different communities. We notice that the 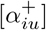 matrices are overall sparse. Some ROIs are not part of any community as they are associated with small coefficients 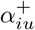. Namely, the messages or representation variance carried by these ROIs are depressed. Thus, it is reasonable to use R-pool to select a few representative ROIs to summarize the group-level representation.

**Fig. 9:**
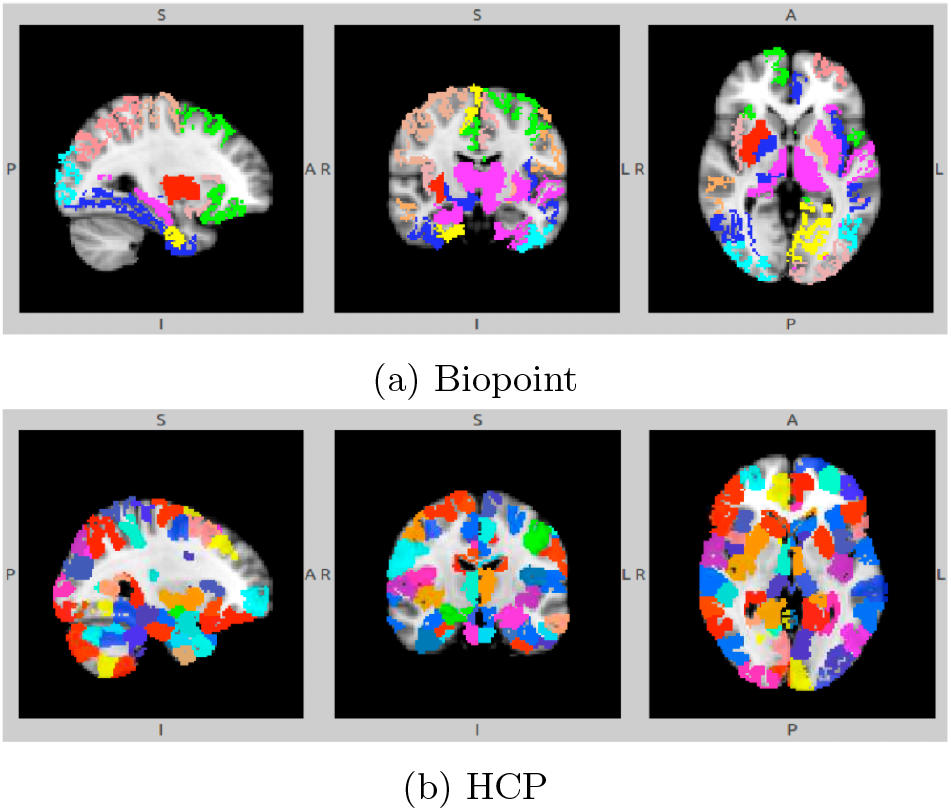
Clustering ROI using 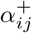 from the 1st Ra-GConv layer. Different colors denote different communities.

**Fig. 10:**
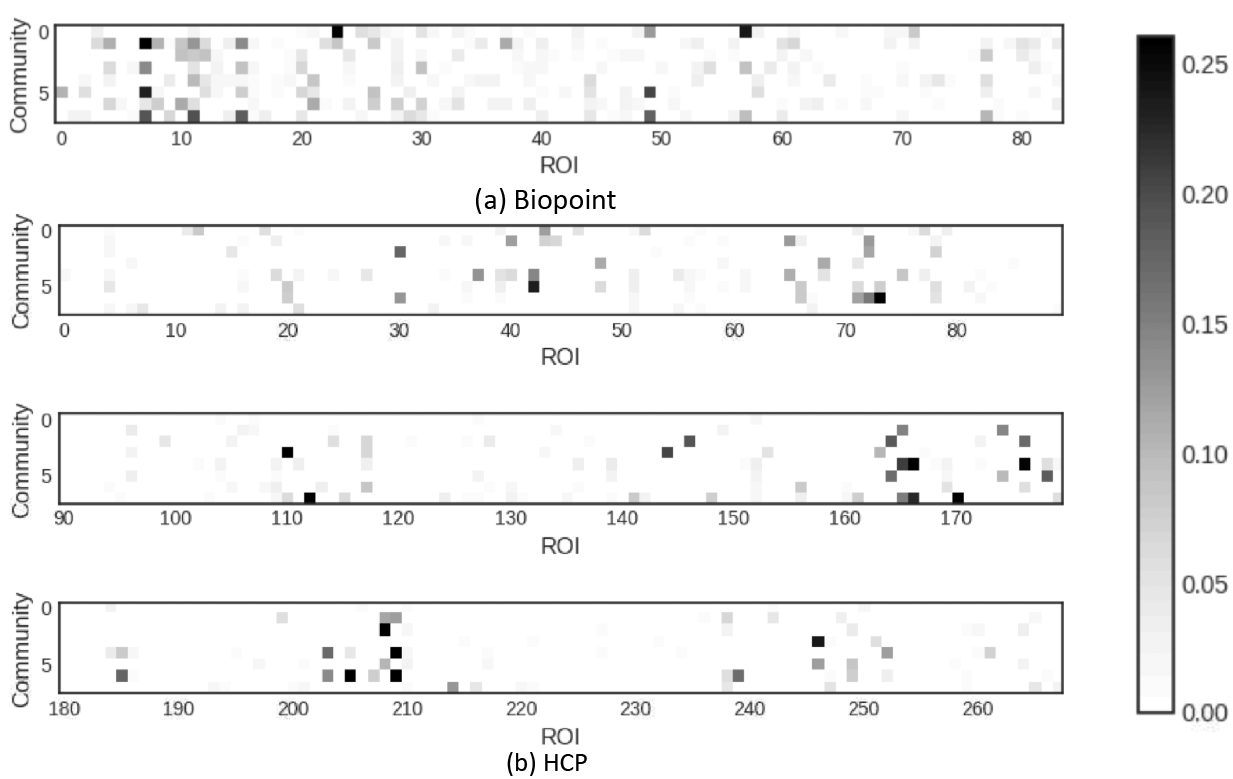
Visualizing Ra-GConv parameter 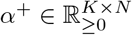, which implies the membership score of an ROI to a community. *K* is the number of communities, represented as the vertical axis. We have *K* = 8 in our experiment. *N* is the number of ROIs, represented as the horizontal axis. (a) is the α^+^ of Biopoint task, and *N* = 84. (b) is the α^+^ of HCP task, and *N* = 268. We split α^+^ of HCP task into three rows for better visualization (note ROI numbering on horizontal axes).

## 4 Discussion

### 4.1 The Model

Our proposed BrainGNN includes (i) novel Ra-GConv layers that efficiently assign each ROI a unique kernel that reflects ROI community patterns, and (ii) novel regularization terms (unit loss, GLC loss and TPK loss) for pooling operations that regulate the model to select salient ROIs. It shows superior prediction accuracy for ASD classification and brain states decoding compared to the alternative machine learning, MLP, CNN and GNN methods. As shown in Fig. 2, BrainGNN improves average accuracy by 3% to 20% for ASD classification on the Biopoint dataset and achieves average accuracy of 94.4% on a seven-states classification task on the HCP dataset.

Despite the high accuracy achieved by deep learning models, a natural question that arises is if the decision making process in deep learning models can be interpretable. From the brain biomarker detection perspective, understanding salient ROIs associated with the prediction is an important approach to finding the biomarkers: the salient ROIs could be candidate biomarkers. Here, we use built-in model interpretability to address the issue of group-level and individual-level biomarker analysis. In contrast, without additional post-processing steps, the existing methods of fMRI analysis can only either perform individual-level or group-level functional biomarker detection. For example, general linear model (GLM), principal component analysis (PCA) and independent component analysis (ICA) are group-based analysis methods. Some deterministic models like connectome-based predictive modeling (CPM) [52,22] (a coarse model averaging edge strengths over entire subject for prediction) and other machine learning based methods provide individual-level analysis. However, model flexibility for different-levels of biomarkers analysis might be required by different users. For precision medicine, individual-level biomarkers are desired for planning targeted treatment, whereas group-level biomarkers are essential for understanding the common characteristic patterns associated with the disease. To fill the gap between group-level and individual-level biomarker analysis, we introduce a tunable regularization term for our graph pooling function. By examining the pairs of inputs and intermediate outputs from the pooling layers, our method can switch freely between individual-level and group-level explanation by end-to-end training. A large regularization parameter for group consistency encourages interpreting common biomarkers for all the instances, while a small regularization parameter allows different interpretations for different instances. However, the appropriate parameters are study-specific and the suitable range can be determined using cross validation. It is worth noting that the individual-level biomarker mentioned in our work is not equivalent to single-subject interpretation, as our methods still require numerous participants for training the model.

### 4.2 Limitation and Future Work

The pre-processing procedure performed in Section 3.1 is one possible way of obtaining graphs from fMRI data, as demonstrated in this work. One meaningful next step is to use more powerful local feature extractors to summarize ROI information. A joint end-to-end training procedure that dynamically extracts graph node features from fMRI data is challenging, but an interesting direction. Also, in the current work, we only try a single atlas for each dataset. For ROI-based analysis, different atlases usually lead to different results [12]. Considering reproducibility and consistency [60,1], it is worth further investigating whether the classification and interpretation results are robust to atlas changes. Although we discussed a few variations of hyperparameters in Section 3.3, more variations should be studied, such as pooling ratio, the number of communities, the number of convolutional layers, and different readout operations. In future work, we will try to understand the interpretation from failure cases and explore how the interpretation results can help improve model performance. We will explore the potential benefits of using BrainGNN to improve GNN-based dynamic brain graph analysis (i.e. [19]). Given the flexibility of GNN to integrate multi-modality data, we will investigate BrainGNN on biomarker detection tasks using an integration of multi-paradigm fMRI data (i.e. [4]). We will explore the connections between the Ra-GConv layers and the tensor decomposition-based clustering methods and the patterns of ROI selection and ROI clustering. For better understanding the algorithm, we aim to work on quantitative evaluations and theoretical studies to explain the experimental results.

## 5 Conclusions

In this paper, we propose BrainGNN, an interpretable graph neural network for fMRI analysis. BrainGNN takes graphs built from neuroimages as inputs, and then outputs prediction results together with interpretation results. We applied BrainGNN on the Biopoint and HCP fMRI datasets. With the built-in inter-pretability, BrainGNN not only performs better on prediction than alternative methods, but also detects salient brain regions associated with predictions and discovers brain community patterns. Overall, our model shows superiority over alternative graph learning and machine learning classification models. By investigating the selected ROIs after R-pool layers, our study reveals the salient ROIs to identify autistic disorders from healthy controls and decodes the salient ROIs associated with certain task stimuli. Certainly, our framework is generalizable to analysis of other neuroimaging modalities. The advantages are essential for developing precision medicine, understanding neurological disorders, and ultimately benefiting neuroimaging research.

## 7 Acknowledgements

Parts of this research was supported by National Institutes of Health (NIH) R01NS035193.

